# A gold standard method for human cell quantification after xenotransplantation

**DOI:** 10.1101/523530

**Authors:** Mona Bensalah, Pierre Klein, Ingo Riederer, Soraya Chaouch, Laura Muraine, Wilson Savino, Gillian Butler-Browne, Capucine Trollet, Vincent Mouly, Anne Bigot, Elisa Negroni

**Affiliations:** Sorbonne Université, Myology Research Center, UM76 and INSERM U974, Institut de Myologie, 75013, Paris, France; Laboratory on Thymus Research, Oswaldo Cruz Institute, Fiocruz, Rio de Janeiro, Brazil; Brazilian National Institute of Science and Technology on Neuroimmunomodulation, Oswaldo Cruz Institute, Oswaldo Cruz Foundation, Rio de Janeiro, Brazil

## Abstract

Xenotransplantation of human cells into immunodeficient mouse models is a very powerful tool and an essential step for the pre-clinical evaluation of therapeutic cell-and gene-based strategies. Here we describe an optimized protocol combining immunofluorescence and real-time quantitative PCR to both quantify and visualize the fate and localization of human myogenic cells after injection in regenerating muscles of immunodeficient mice. Whereas real-time quantitative PCR-based method provides an accurate quantification of human cells, it does not document their specific localization. The addition of an immunofluorescence approach using human-specific antibodies recognizing engrafted human cells gives information on the localization of the human cells within the host muscle fibres, in the stem cell niche or in the interstitial space.

These two combined approaches offer an accurate evaluation of human engraftment including cell number and localization and should provide a gold standard to compare results obtained either using different types of human stem cells or comparing healthy and pathological muscle stem cells between different research laboratories worldwide.

## Introduction

Investigating the *in vivo* behaviour of human cells, whether it concerns fundamental aspects, pathological mechanisms or therapeutic strategies, represents a challenging aspect of cell biology. To recreate *in vivo* dynamics and provide knowledge about *in vivo* human cell behaviour, xenotransplantation of human cells in immunodeficient mice has been developed and represents a unique experimental approach. Although xenotransplantation has been widely used in immunology, it also represents a powerful tool to investigate the cell mechanisms involved in both skeletal muscle repair and homeostasis in normal and pathological situations [1]. Post-mitotic and stable during healthy adulthood, skeletal muscle can face rapid and devastating changes following trauma or in dystrophic conditions [1]. At rest, the physiological muscle stem cells, called satellite cells, are mitotically quiescent. After injury or increased load, satellite cells become activated and proliferate thus supplying a large number of new myonuclei ready to fuse to replace damaged myofibers and assure a rapid muscle growth and repair, in addition to generating new satellite cells by self-renewal [2]. Muscle regeneration is an extremely well orchestrated process in which several cell actors (satellite-, immune system-and non-myogenic-cells) [3-5] play distinct roles at different steps of muscle regeneration. Slight modifications in the regeneration kinetics can result in impaired muscle repair. In healthy conditions, muscle regeneration is highly efficient whereas in muscular diseases, regeneration can be delayed or compromised, resulting in progressive muscle wasting and weakness. To better understand muscle maintenance and repair in healthy and dystrophic contexts, as well as to evaluate the efficacy of cell-based therapeutic strategies, it is essential to study the *in vivo* behaviour of control, dystrophic-derived, and/or modified human satellite cells. Indeed, for genetic diseases including muscular dystrophies, cell-based treatment is one of the innovative and promising therapeutic approaches [6]. In addition, genetic modifications of human stem cells, whether it is through gene therapy or direct genomic correction, must be tested in an *in vivo* context as a necessary step towards therapy. In this context, accurate methods are needed to evaluate human cell behaviour and participation to muscle regeneration during xenotransplantation. Different cell types have been used as possible vectors for cell-based therapy [7, 8, 9, 10, 11]. However, the direct comparison of distinct cell types or treatments is often difficult since different techniques are being used in different laboratories to quantify *in vivo* engraftment. Here we describe a combined immunofluorescence and real time quantitative PCR-based approach to quantify the grafting efficacy of human myogenic precursors after intramuscular injection in immunodeficient mouse and to analyse their fate in these regenerating muscles.

## Materials and methods

### Animals

2-3 month old Rag2^-/-^Il2rb^-/-^ immunodeficient mice were used as recipients for human myogenic precursor transplantation. Mice were anaesthetized by an intraperitoneal injection of 80 mg/kg of ketamine hydrochloride and 10 mg/kg xylasine (Sigma-Aldrich. St. Louis, MO) as previously described [12]. This study was carried out in strict accordance with the legal regulations in France and according to European Union ethical guidelines for animal research. The protocol was approved by the Committee on the Ethics of Animal Experiments Charles Darwin N°5 (Protocol Number: 7082-2016092913021452). All surgery was performed under ketamine hydrochloride and xylasine anesthesia, and all efforts were made to minimize suffering.

### Cultures of human myoblasts

Human myoblasts were isolated from the quadriceps muscle of a 5-day-old infant in accordance with the French legislation on ethical rules, as previously described [13]. Cells were expanded in a growth medium consisting of 199 medium and DMEM (Invitrogen, Life Technologies, Saint-Aubin, France) in a 1:4 ratio, supplemented with 20% foetal calf serum and 50 μg/ml gentamycine (Invitrogen), at 37°C in a humid atmosphere containing 5% CO2. Population doublings (PDs) were determined at each passage according to the formula: log (N/n)/log 2 where N is the number of cells counted and n is the number of cells initially plated.

### Cell preparation and myoblast transplantation

Myoblast cultures were washed in PBS, trypsinized, centrifuged, and re-suspended in PBS. The cells were injected into both *Tibialis Anterior* (TA) muscles. Prior to myoblast implantation, the TAs of the immunodeficient mice were subjected to three freeze lesion cycles of ten seconds each in order to damage the muscle fibers, and trigger regeneration, thus stimulating the implanted myoblasts to fuse and form new muscle fibers. Human myoblasts were implanted into the recipient’s muscle immediately after cryodamage, using a 25 μl Hamilton syringe as previously described [7]. 15 μl of cell suspensions containing 5 × 10^5^ myoblasts in PBS were injected in a single injection site in the mid belly of the TA. For combined immunofluorescence (IF)/qPCR analysis 1 × 10^5^ cells were injected. The skin was then closed using fine sutures. At 0, 5, 7, 15, 21, 30 days (d) after injection mice were sacrificed (3≤n≤6 for each time point) and the TAs were dissected and analysed. TAs were mounted in tragacanth gum (6% in water; Sigma-Aldrich, St Louis, MO) and frozen in isopentane precooled in liquid nitrogen for immunofluorescence (IF) analysis and combined IF/qPCR analysis. TAs were directly frozen in liquid nitrogen for DNA extraction and qPCR analysis. All muscles were stored at −80°C.

### Histology and immunofluorescence

For the assessment of tissue morphology and analysis at different steps of muscle regeneration, 5 μm thick muscle cryosections were stained with hematoxylin and eosin (H&E) and examined by light microscopy. Immunofluorescence analyses of grafted TA muscles were performed using antibodies specific for: human lamin A/C (mouse monoclonal IgG2b, 1:400, NCL-Lam-A/C, Novocastra, Leica Biosystems, Wetzlar, Germany or mouse monoclonal IgG1, 1:400, clone JOL2, AbCam, Cambridge, UK) to detect human nuclei; Pax7 (mouse monoclonal IgG1, 1:50, Pax7 Developmental Studies Hybridoma Bank) to detect satellite cells; human spectrin (mouse monoclonal IgG2b, 1:50, NCL-Spec1, Novocastra) to detect human protein expressing fibres; laminin, (1:400, rabbit polyclonal, Z0097, Dako) to delineate muscle fibre architecture. Briefly, cryostat sections were blocked in 2% BSA in PBS. Sections were then incubated with primary antibodies for 1h, washed in PBS, and subjected to the appropriate secondary antibodies for 45 min: Alexa Fluor 488 coupled goat anti-mouse IgG2b, Alexa Fluor 555 coupled goat anti-mouse IgG1 and Alexa Fluor 647 coupled goat anti-rabbit. Hoechst staining (0.5μg/ml, Hoechst No. 33258; Sigma-Aldrich) was used to visualize nuclei and sections were mounted using an aqueous medium (Cytomation fluorescent mounting Medium, S3023, Dako). All images were visualized using an Olympus BX60 microscope (Olympus Optical, Hamburg, Germany), digitalized using the Photometrics CoolSNAP fx CCD camera (Roper Scientific, Tucson, AZ) and analyzed using the MetaView image analysis software (Universal Imaging, Downington, PA). To calculate nuclear size, a total of at least 500 cells were counted in different cross-sections at 7 days after injection using Image J 1.44o analysis software (http://imagej.nih.gov/ij).

### Isolation of genomic DNA from injected TA

Genomic DNA was extracted using a salting-out procedure. Briefly, TAs were resuspended in proteinase K digestion buffer (50mM Tris pH 7,4, 20mM EDTA, 1% SDS) containing 0,2 mg/ml proteinase K. To obtain a lysis of the muscle tissue, FastPrep system (MP Biomedicals, France) was used to completely homogenize the injected TAs. Muscle homogenization was incubated at 37°C overnight under agitation. After proteinase K inactivation and RNase A treatment (1 mg/ml), saturated 6M NaCl solution was added to each tube and shaken vigorously for 30 seconds, followed by centrifugation at 12000 rpm for 7 minutes (min). In order to precipitate the DNA, 2 volumes of absolute ethanol were added to the supernatant. After centrifugation (5000g for 5 min), DNA pellet was resuspended in ethanol at 70%. After centrifugation (5000g for 5 min), DNA was allowed to dissolve in the same volume of water over night at 4°C before quantification. DNA was quantified using ND-1000 NanoDrop spectrophotometer (Thermo Fischer Scientific, DE, USA). To establish a standard curve, increasing quantity of human myoblasts, ranging from 5 × 10^3^ to 5 × 10^5^, was added to a mouse TA in an eppendorf tube and frozen in liquid nitrogen. DNA extraction was then performed as described above.

### Quantitative-PCR

Quantitative polymerase chain reaction (qPCR) was carried out using SYBR green mix buffer (Roche Applied Science, Meylan, France) in a LightCycler 480 Real-Time PCR System (Roche Applied Science) as follows: 8 min at 95°C followed by 50 cycles at 95°C for 15 seconds (s), 60°C for 15 s and 72°C for 15 s, with the program ending in 5 s at 95°C and 1 min at 65°C. Specificity of the PCR product was checked by melting curve analysis using the following program: 65°C increasing by 0.11°C/second to 97°C. Primer sequences for human Titin (TTN) DNA, designed using the web interface Primer-BLAST http://www.ncbi.nlm.nih.gov/tools/primer-blast/, are: 5’-ACCACATGCATTTTATCAGAGC-3’ and 5’-CCTTGAACATTCTCAACAGGC-3’.

### Statistical Analysis

Data were expressed as the mean  SEM. All statistical analyses were performed using GraphPad Prism (version 6.0d, GraphPad Software Inc., San Diego, CA). Statistical significance was assessed by one-way ANOVA or student *t*-test. A difference was considered to be significant at \**P* < 0.05, \*\**P* < 0.01, \*\*\**P* < 0.001.

## Results and discussion

### Real time PCR quantification of human engrafted cells: design and specificity of primers

Following muscle injury, satellite cells, which are the muscle stem cells, become activated, start to proliferate and fuse together to replace damaged myofibers. In order to create a regenerative environment in the host and follow the participation of human satellite cells to tissue repair, we induced regeneration in the Tibialis Anterior (TA) muscle of immunodeficient mice using a freeze-injury protocol [7], one of the most commonly used physical procedures for provoking muscle injury and repair in mice [15]. Human myoblasts were injected, after cryodamage, into regenerating TA and at 0, 5, 7, 15, 21 or 30 days (d) after injection, mice were sacrificed, the TAs excised and processed (Fig 1A).

**Fig 1.**
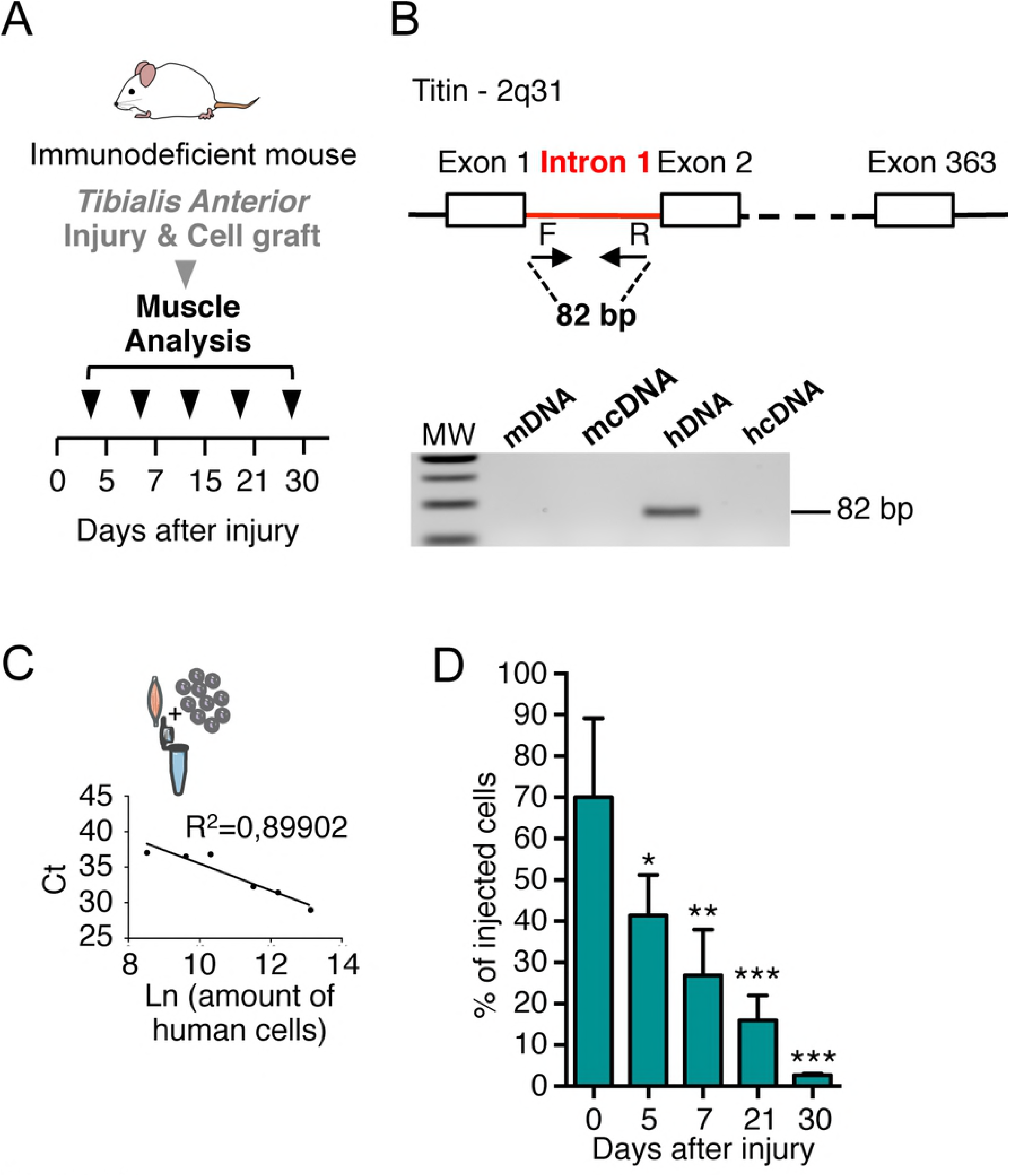
Quantification of human myoblasts in TA muscle by qPCR analysis. (**A**) Experimental scheme of the kinetic of analysis of TA muscles injected with human myoblasts after freeze-injury. Human myoblasts were injected in a single injection site in the mid belly of the TA after that an injury was produced on the TA. TA muscles were dissected and frozen at different time points (0, 5, 7, 15, 21 or 30 days) after cell injection. (**B**) Schematic representation of primer (arrows) used to amplify human titin DNA by qPCR. Species specificity of titin primers is shown using mouse DNA/cDNA and human DNA/cDNA by qPCR and migration on agarose gel. (**C**) Representative standard curve obtained with DNA mouse TA muscles mixed and extracted with different amount of human cells. In the graph the number of cells is expressed as the Ln of the number of cells. In the ordinate axis the Ct (cycle threshold) representative of a qPCR run are reported. (**D**) Histograms showing the rapid quantification using qPCR analysis of TA analysed at 0, 5, 7, 21 and 30d after injection of human cells. The number of human myoblasts was calculated from the qPCR calibration curve. Results are means of the different TAs analysed ± SEM. The mean of each column is compared with the mean of the 0d column using an ordinary one-way ANOVA followed by Dunnett’s multiple comparisons test. \**P* < 0.05, \*\**P* < 0.01, \*\*\**P* < 0.001.

In order to estimate the number of engrafted human myoblasts present in the TA after injection, we quantified the amount of human DNA present in the mouse muscles, using primers recognizing specifically the human Titin (*TTN*) gene (and not mouse *ttn*). The human gene encoding for *TTN* (367 exons) is found as only one single copy, localized on chromosome 2 (2q31). To ensure that our primers recognize exclusively genomic DNA sequences, we designed the couple of primers in the intron 1 of the human gene. Specificity of the primers has been assessed by PCR and results are shown in Fig 1B.

In order to establish a standard curve for the quantification by qPCR of the amount of human cells, we prepared a range of samples by mixing a defined number of human myoblasts (5 × 10^3^, 15 × 10^3^, 3 × 10^4^, 1 × 10^5^, 2 × 10^5^, 5 × 10^5^ cells) together with an immunodeficient mouse TA previously frozen in liquid nitrogen. This procedure allows us to mimic the DNA extraction of human cells after injection in mouse TA, maintaining the same background, that is to say the presence of a large quantity of mouse DNA and contaminants accumulated during the DNA extraction, potentially inhibiting the qPCR (Fig 1C). In order to easily compare standard curve samples and research samples the same experimental process was applied to both: same dilution of the total extracted DNA, and same volume for each sample in each 384-plate well for qPCR analysis. We found a linear correlation with R^2^=0,89902 between the amount of cells (expressed as Ln) and the cycle threshold (Ct) value (Fig 1C).

### qPCR for human engrafted cells quantification

The total DNA extracted was resuspended in distilled water and an aliquot of diluted extracted DNA was used to perform the qPCR. The qPCR technique, performed independently on 4 TAs and reproduced in triplicate, allows a rapid quantification of a large number of samples. Analysis of the qPCR data showed a progressive diminution of the number of human myoblasts from 5d after grafting when compared to the amount found at injection (0d) (Fig 1D). This is in accordance with what is found in our and other previous works with human cells [12, 16-17]: human myoblasts persist better than murine cells in mouse muscles and the massive loss of murine cells observed during the first days after injection [18] is not confirmed with human cells, when they are injected in an immunodeficient mouse, although a loss of cells is gradually observed at later time points (Fig 1C). More importantly, immediately after injection, human myoblasts remain confined in an enriched laminin “pocket” around the injection site [12]. In this pocket, the cells are tightly packed together making their quantification by immunofluorescence analysis very difficult and time-consuming. qPCR analysis, which easily permits quantification at these time points, has been used to quantify the amount of human DNA present in the samples. Our results confirm that the qPCR–based approach represents a robust and sensitive technique to detect human DNA in a murine context, as recently also described by Funakoshi et al, with qPCR probes specific for human Alu elements in mouse DNA [19].

### Immunofluorescence analysis: spatial information

In order to visualize grafted human cells, TA were entirely cut along their entire length into 5-μm thick cryosections. A schematic representation of the analysis of TA muscles injected with human myoblasts after freeze-injury and the method used to analyse the TAs is given in Fig 2A. Briefly serial sections were collected on microscope slides (n=30), each section being spaced from the next section by 500 μm (Fig 2A). One slide was used to quantify the total number of human cells over the entire length. Human nuclei were revealed using a human-specific anti lamin A/C antibody. This immunostaining allows the localization of human cells at early time points, or myonuclei and human undifferentiated cells at later time points, within the injected muscle, and thus the global dispersion on transversal sections. This dispersion can be evaluated by measuring the area occupied by the injected cells as previously shown [7] using imaging software. Analysis was carried out on the section bearing the largest number of lamin A/C-positive cells along the entire muscle. On the transversal sections, the smallest ellipse area containing inside of it all of the human cells was calculated as a percentage of the total area of the TA using Image J analysis system software. We chose to analyse this parameter 21d after injection, since this time point is the one that has been most often used to analyse the participation of myogenic cells to muscle regeneration. At this time, human myoblasts are found in an area representing 50,74% ±5.5 of the whole transversal section of the muscle (Fig 2B). Longitudinal dispersion of injected cells within the host muscle can be evaluated by measuring the distance (*i.e.*, number of sections) along which the presence of lamin A/C positive nuclei are detected throughout the entire longitudinal length of the muscle. This analysis showed that at 21d after injection, human cells are present throughout roughly 70% (70,23±4,9) of the length of the injected muscle (Fig 2B). H&E staining was performed to appreciate the regeneration status and the architecture of the damaged muscle: typically a control muscle appears well organized with mouse myofibers characterized by a polygonal shape and peripheral nuclei (Fig 2C, CTR). 7d after cryodamage, we observe an intense muscle regeneration, clearly identified by multicentronucleated small fibres. Many inflammatory cells were still present in the interstitial space of the tissue. At 15d after cryodamage, almost all mouse fibres were centrally nucleated. Muscle fibres recovered their original size from 21d after injury, although all of the regenerated fibres still had centrally located nuclei (Fig 2C). Anti-Lamin A/C antibody (human specific) together with an anti-laminin antibody recognizing mouse and human laminin allowed the visualisation of human cells within the mouse muscle either inside murine fibres as myonuclei or between muscle fibres in an interstitial position (Fig 2C and 2D). A co-staining (Fig 2E) using an anti-lamin AC (staining nuclear envelope), anti-laminin (staining the basal lamin) and anti-Pax7, a marker of satellite cells, show that human cells can colonise the tissue specific stem cell niche, thus amplifying their therapeutic potential during successive cycles of regeneration, as it occurs in muscle dystrophies. The ability to colonize the satellite cell niche can vary depending on the myogenic cell type. As an example, we and other groups showed that different types of human muscle stem cells (myoblasts derived satellite cells, pericytes, CD133 expressing cells) contribute in different proportions to the satellite cell pool *in vivo* when injected intramuscularly [6]. In the cryodamaged mouse model presented here, 21d after injection, 5,76% of human injected myoblasts were found in the satellite cell position, 38,90% were found inside the myofibers as myonuclei and 55,33% were located in the interstitial spaces, outside muscle fibres (Fig 2E). Finally, a double staining using human specific antibodies anti lamin AC/dystrophin or lamin AC/spectrin (Fig 2F) allowed us to detect the formation of chimeric fibres expressing the human proteins and therefore the participation of human cells to fibre regeneration, which reflects their therapeutic potential to bring a missing protein in a dystrophic context.

**Fig 2.**
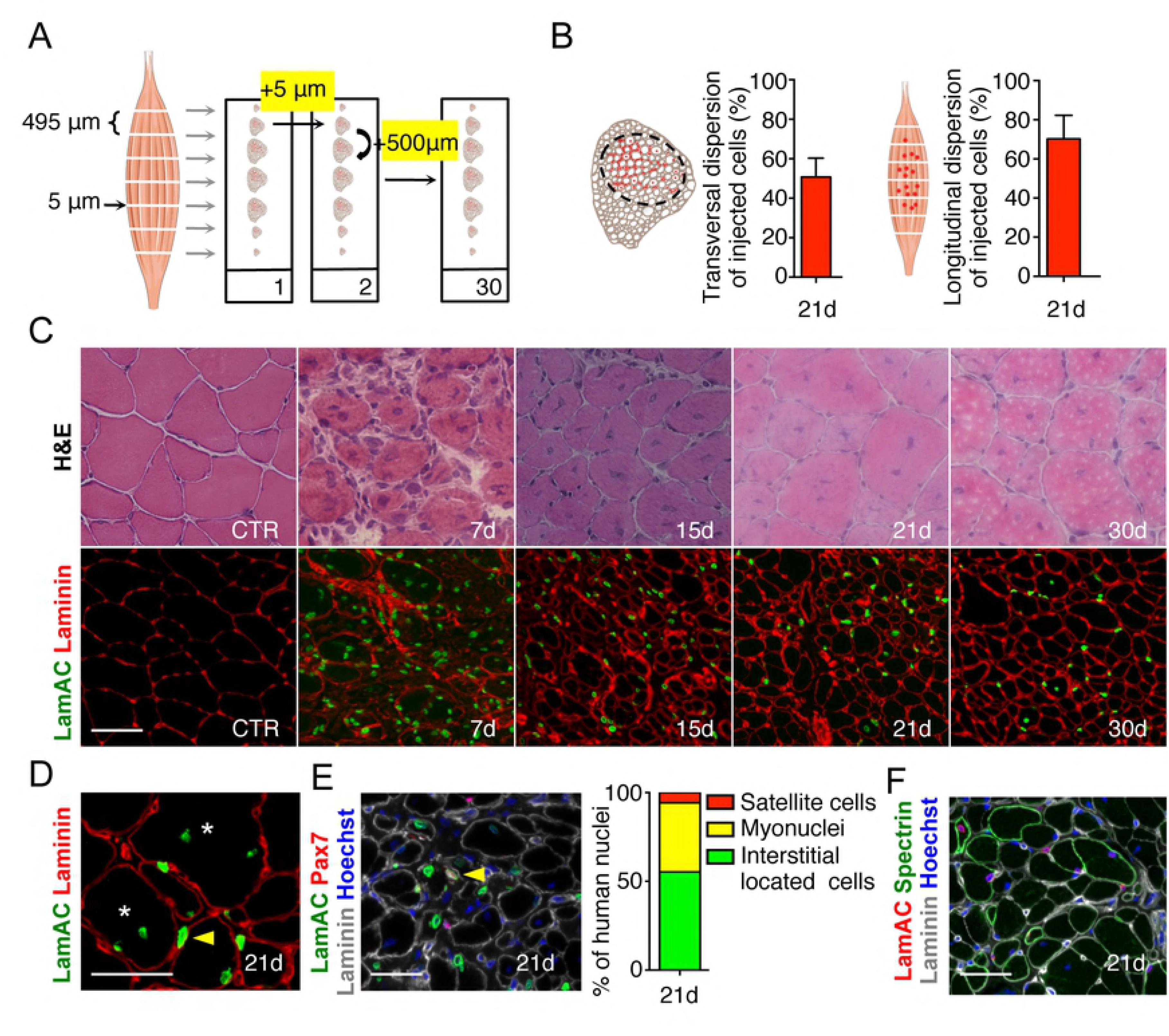
Identification of human myoblasts in TA muscle by immunofluorescence. (**A**) Schematic representation of the immunofluorescence (IF) processing of a TA muscle. Each section (5 μm) on the same slide is separated from the previous and from the following one by 500 μm. (**B**) Representative transversal mouse section and quantification of the transversal dispersion of human injected cells 21d after injury and injection. Schematic longitudinal representation of a TA muscle and quantification of the longitudinal dispersion of human injected cells 21d after injury and injection. (**C**) Representative H&E transverse sections of control TA muscle and 7, 15, 21 and 30d after freeze-injury and IF showing injected human cells. Human cells are visualised using a human specific lamin AC antibody (LamAC, green) and the extra cellular laminin protein is visualised in red. (**D**) Human myoblasts can be found as myonuclei (asterisk) or in the interstitial space (yellow arrow point). Human myoblasts are detected with an antibody raised against human lamin AC (green). Laminin is visualised in red. (**E**) Immunostaining using antibodies directed against human LamAC (green), Pax7 (red) and laminin (grey). Nuclei are counterstained with Hoechst (blue). Human LamAC/Pax7 satellite cells (yellow arrows) are localized under the basal lamina (grey). Histogram showing the quantification of the *in vivo* repartition of human myoblasts 21d after injection. All LamAC positive cells are counted and classified according to their position and the marker they express. (**F**) Representative transverse section showing human myoblasts participation to mouse muscle regeneration. Human nuclei are revealed using a human-specific anti-LamAC antibody (red), and chimeric fibres expressing human proteins are visualized using an anti-human spectrin–specific antibody (green). An anti-laminin antibody is used to delineate muscle structure (grey). Nuclei are counterstained with Hoechst (blue) Scale bar = 50 μm.

### Estimation of the number of engrafted human cells

We propose a method to quantify the total number of human cells persisting in the TA, taking into account the size of the injected cells and the size of the cryosections (s=5μm). We measured the size of the nucleus, using the lamin A/C staining that we used to identify human myoblasts in the mouse muscle (Fig 2B and 2C). To calculate the average nuclear diameter (D), a total of at least 500 nuclei were counted 7d after injection using Image J software. Nuclear diameter at 7d post-injection was 8,0 ± 2,5 μm (Fig 3A). The linear density (n) of cells can be obtained by dividing the number of cells (N) counted in a section over the section size s (5μm). But given its finite size D, each cell could be visible in a number of sections. The linear density (n) should, therefore, be weighted by the average number of sections over which each cell is detected [(D+s)/s], as follows:

**Fig 3.**
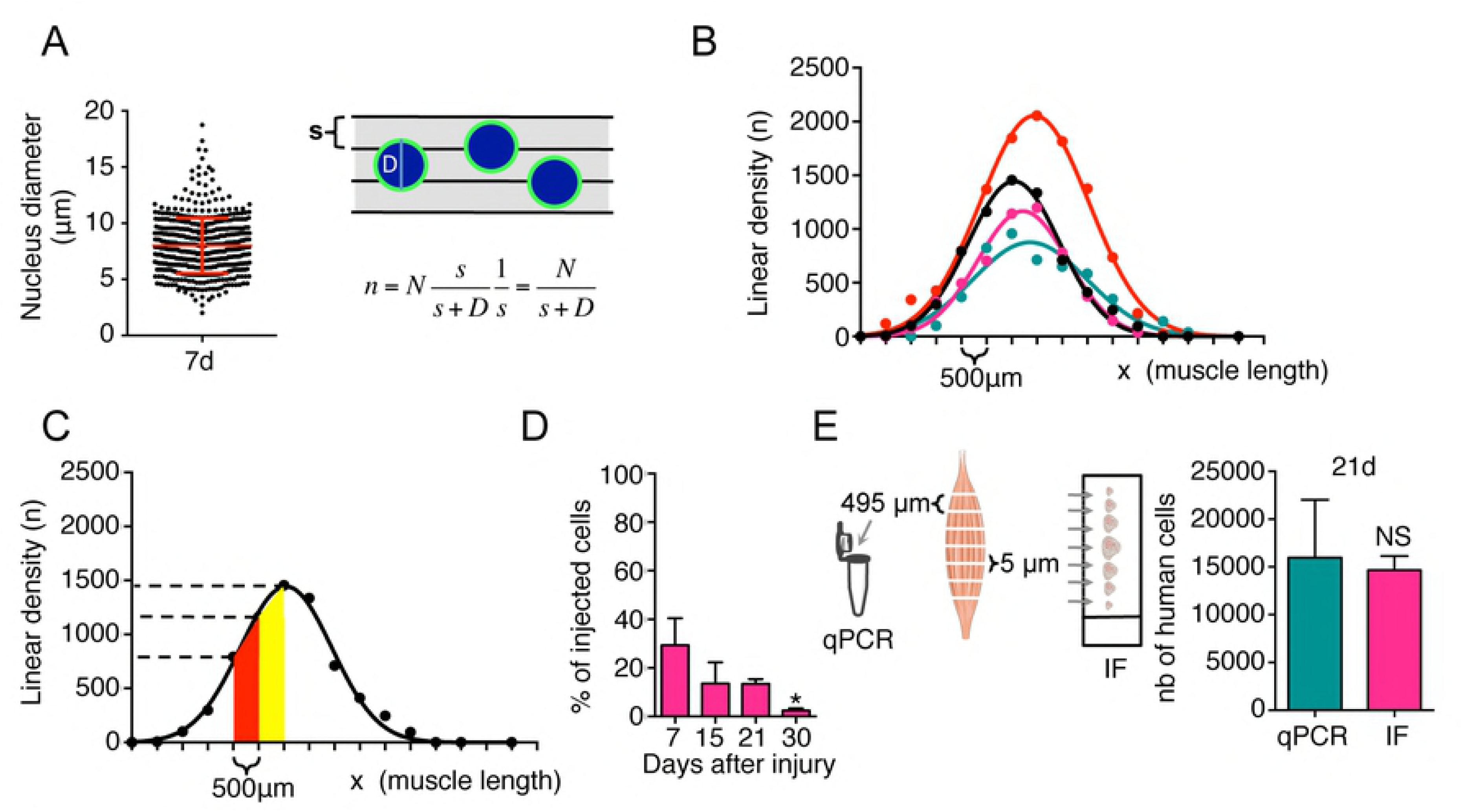
Quantification of human myoblasts in TA muscle by immunofluorescence method. (**A**) Size of human myoblast nuclei 7 d after injection *in vivo.* At least 500 nuclei were counted in different cross-sectional sections. Schematic representation of nuclei in the muscle sections and linear density (n) of the nuclei in the muscle length. s= size of muscle section; D= nucleus diameter; n=linear density; N= number of cells counted in a muscle section (**B**) Example of typical curves obtained counting human myoblasts 30 d after a single injection in the middle of the TA. Curves show the linear density of 4 independent injections. (**C**) Graph showing an example of the linear density (n) of human cells in the muscle. (**D**) Quantification of human cells found 7, 15, 21 and 30d after grafting in cryodamaged muscle using the IF analysis. Results are means of the different TAs analysed ± SEM. The mean of each column is compared with the mean of the 7d column using an ordinary one-way ANOVA followed by Dunnett’s multiple comparisons test. \**P* < 0.05, \*\**P* < 0.01, \*\*\**P* < 0.001. (**E**) Quantification of human myoblasts by quantitative PCR and IF methods. Schematic representation of the two processing methods and histograms showing values obtained with the two techniques 21d after freeze-injury and injection of human myoblasts. Results are means of the different TAs analysed ± SEM. Student *t*-test was used to compare differences between two groups. NS: non significant).

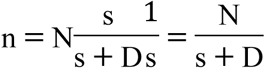

To estimate the total number of cells one has to integrate the linear density n of the cells over the entire length (x) of the muscle (Fig 3B and 3C).

We counted cells only in 5μm sections separated by 500μm (L). The number of cells in the intermediate sections was estimated by linear interpolation (Fig 3C), and numerical integration was performed using the formula, given below:

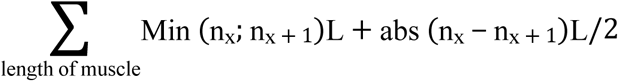

where Min represents the minimum value between n_x_ and n_x+1_ and abs the absolute value. Using this approach, we found a diminution in the number of human cells from d7 to d30 post injection, which was consistent with the qPCR analysis (Fig 3D).

### Combinative analysis of muscle samples: estimation of the number of engrafted human cells by qPCR and immunofluorescence

In order to compare the two methods, we processed the same TA muscle sample at 21 d after injection of 1 × 10^5^ human myoblasts, using both techniques in parallel (Fig 3E). Interspaced sections were processed for immunofluorescence analysis as described above, and intermediate slides were used for qPCR analysis. Both quantifications of human cells show no statistical difference between the two methods (IF: 14657±857, qPCR: 15966±3507, NS), indicating that both methods are equally efficient quantitative tools (Fig 3E).

## CONCLUSIONS

In the present study, we have evaluated two different protocols for the quantification of human cells after transplantation into a muscle of immunodeficient mice. Our results show that both of these methods are equally accurate to quantify human cells within the recipient mouse muscle since results obtained with these two methods are not significantly different (Fig 3E). Each method has its specificity: the qPCR-based method can be used for a quantitative, precise and rapid estimation of human material in human xenografts, but does not give any information about the localization of the engrafted cells and modifications of the architecture of the host tissue. Immunofluorescence, on the other hand, provides spatial information but is more time consuming regarding the quantification of human material. Most importantly, we show that both methods can be combined using the same starting material: a few cryosections can be processed by immunofluorescence to check the localization of human nuclei, and the fate and behaviour of the injected cells while qPCR will provide rapidly quantitative data. We propose the combination of these two methods as a gold standard to compare results obtained with different types of human myogenic stem cells (different precursors, control or pathological, treated or not, *etc.)* or when different mouse models are used. Such combined methods can be used to assess results obtained in different laboratories, often difficult to compare, in a more objective and rigorous way. Lastly, we believe that the combination of these two approaches can be applied to analyse more generally grafting and cell transplantation experiments, independently of the target tissue and that they may contribute to a variety of transplantation studies aimed at analysing human cells in mouse models.

## Acknowledgments

We thank Benoît Darquié and Athanasios Laliotis for helpful discussion and suggestions. This work has been supported by grants from the AFM (Association Française contre les Myopathies, OPMD Network Research Program 17110), INSERM and Sorbonne Université (EMERGENCE 2016, SU-16-R-EMR-60), CNPq and Faperj (Brazil). We thank Maud Chapart and Stéphane Vasseur of MYOBANK-AFM (BB-0033-00012).

